## Supplemental Information for "Cell motility modes are selected by the interplay of mechanosensitive adhesion and membrane tension"

May 31, 2023

#### 1 Linear stability analysis

We provide details of the linear stability analysis of the governing equations, with the goal of deriving the dispersion relations of Eqs. [9] and [10] of the main text. The dependent variables in the problem are the shape $R(\theta,t)$ of the cell, the local off-rate $r(\theta,t)$, the local fraction of bound linker $n(\theta,t)$, and the membrane tension $\sigma_m(t)$. In the base state, denoted by overbars, the cell is circular with radius 1 ($\bar{R}=1$), and the retrograde velocity balances the polymerization velocity everywhere: $v_p = \bar{r}\log\bar{r}$. The fraction of bound linkers is uniformly distributed along the cell edge with value $\bar{n} = r_{\text{on}}/(r_{\text{on}}+\bar{r})$, and the force balance provides the membrane tension as

$$\bar{\sigma}_m = \frac{1}{1+h/\bar{R}}(\bar{r}\log\bar{r} + \zeta_1\bar{n}\log\bar{r} - \sigma_c). \tag{1}$$

The variables are perturbed as

$$R(\theta,t) = \bar{R} + \delta R(\theta,t), \quad r(\theta,t) = \bar{r} + \delta r(\theta,t), \quad n(\theta,t) = \bar{n} + \delta n(\theta,t), \quad \sigma_m(t) = \bar{\sigma}_m + \delta\sigma_m(t), \tag{2}$$

where the perturbations are assumed to be small in magnitude. The local curvature of the cell edge $\kappa$ can be obtained from the shape, and, to leading order, is expressed as

$$\kappa \approx \frac{1}{\bar{R}} - \frac{\delta R}{\bar{R}^2} - \frac{1}{\bar{R}^2}\frac{\partial^2 \delta R}{\partial \theta^2}. \tag{3}$$

Linearizing the governing equations [7] of the main text yields the system

$$\frac{\partial \delta R}{\partial t} = -\epsilon(1+\log\bar{r})\delta r, \tag{4a}$$

$$\frac{\partial \delta n}{\partial t} = -(r_{\text{on}}+\bar{r})\delta n - \bar{n}\delta r + \frac{D}{\bar{R}^2}\delta n_{\theta\theta}, \tag{4b}$$

$$\frac{\mathrm{d}\delta\sigma_m}{\mathrm{d}t} = -k_\sigma \int_0^{2\pi} (1+\log\bar{r})\bar{R}\,\delta r\,\mathrm{d}\theta, \tag{4c}$$

$$-\frac{h\bar{\sigma}_m}{\bar{R}^2}(\delta R + \delta R_{\theta\theta}) + \Big(1+\frac{h}{\bar{R}}\Big)\delta\bar{\sigma}_m = \Big(1+\log\bar{r}+\zeta_1\frac{\bar{n}}{\bar{r}}\Big)\delta r + \zeta_1\log\bar{r}\,\delta n. \tag{4d}$$

We assume normal modes for the perturbation variables,

$$\delta R = \sum_{k=0}^{\infty} R_k \exp(\mathrm{i}k\theta + \lambda_k t) + c.c., \qquad \delta r = \sum_{k=0}^{\infty} r_k \exp(\mathrm{i}k\theta + \lambda_k t) + c.c., \tag{5}$$

$$\delta n = \sum_{k=0}^{\infty} n_k \exp(\mathrm{i}k\theta + \lambda_k t) + c.c., \qquad \delta \sigma_m = \sum_{k=0}^{\infty} \sigma_k \exp(\lambda_k t) + c.c., \tag{6}$$

where $k \in \mathbb{N}$ are integer wavenumbers, $\lambda_k$ are the corresponding growth rates, and *c.c.* denotes the complex conjugate. Inserting these expressions into Eqs. [4] provides

$$\lambda_k R_k = -\epsilon(1+\log\bar{r})r_k, \tag{7a}$$

$$\lambda_k n_k = -(r_{\mathrm{on}}+\bar{r})n_k - \bar{n}r_k - \frac{D}{\bar{R}^2}k^2 n_k, \tag{7b}$$

$$\lambda_k \sigma_k = -2\pi\delta_{k0}k_\sigma(1+\log\bar{r})\bar{R}\,r_k, \tag{7c}$$

$$-\frac{h\bar{\sigma}_m}{\bar{R}^2}(1-k^2)R_k + \Big(1+\frac{h}{\bar{R}}\Big)\sigma_k = \Big(1+\log\bar{r}+\zeta_1\frac{\bar{n}}{\bar{r}}\Big)r_k + \zeta_1\log\bar{r}\,n_k. \tag{7d}$$

When deriving Eq. [7c], we have used that $\int_0^{2\pi}\exp(\mathrm{i}k\theta)\,\mathrm{d}\theta = 2\pi\delta_{k0}$ where $\delta_{k0}$ is the Kronecker delta. In particular, this shows that the tension perturbation at most involves mode $k=0$, consistent with the fact that the tension is independent of $\theta$: $\delta\sigma_m = \sigma_0\exp(\lambda_0 t) + c.c.$, and $\sigma_k = 0$ for $k \geq 1$. Given this observation, modes $k=0$ and $k \geq 1$ require slightly different treatment and are analyzed separately.

#### Mode $k = 0$

Eliminating the growth rates $\lambda_0$ in Eqs. [7] when $k=0$ leads the following constraints on the modes amplitudes:

$$r_0 = \frac{r_{\mathrm{on}}+\bar{r}}{\epsilon(1+\log\bar{r})/R_0 - \bar{n}/n_0}, \qquad \sigma_0 = \frac{2\pi\bar{R}k_\sigma}{\epsilon}R_0, \tag{8a}$$

$$\bar{n}LR_0^2 + \Big[\Big(1+\log\bar{r}+\zeta_1\frac{\bar{n}}{\bar{r}}\Big)(r_{\mathrm{on}}+\bar{r}) - \epsilon(1+\log\bar{r})L - \zeta_1\bar{n}\log\bar{r}\Big]n_0R_0 + \epsilon\zeta_1\log\bar{r}(1+\log\bar{r})n_0^2 = 0, \tag{8b}$$

where we have defined

$$L = \frac{2\pi\bar{R}k_\sigma}{\epsilon}\Big(1+\frac{h}{\bar{R}}\Big) - \frac{h\bar{\sigma}_m}{\bar{R}^2}. \tag{9}$$

Equation [8b] is a quadratic equation for $R_0$ when $n_0$ is specified. The other amplitudes $r_0$ and $\sigma_0$ can then be obtained using Eq. [8a]. The two solutions corresponds to two independent modes of fluctuation around the base state.

Eliminating the initial amplitudes of perturbations then provides the eigenvalue equation for the growth rate for the $k = 0$ mode:

$$\left(1+\log\bar{r}+\zeta_1\frac{\bar{n}}{\bar{r}}\right)\lambda_0^2+\left\{\left(1+\log\bar{r}+\zeta_1\frac{\bar{n}}{\bar{r}}\right)(r_{\rm on}+\bar{r})+\left[\left(1+\frac{h}{\bar{R}}\right)2\pi\bar{R}k_\sigma-\frac{\epsilon h\bar{\sigma}_m}{\bar{R}^2}\right](1+\log\bar{r})-\zeta_1\bar{n}\log\bar{r}\right\}\lambda_0$$
$$+\left[\left(1+\frac{h}{\bar{R}}\right)2\pi\bar{R}k_\sigma-\frac{\epsilon h\bar{\sigma}_m}{\bar{R}^2}\right](1+\log\bar{r})(r_{\rm on}+\bar{r})=0. \tag{10}$$

#### Modes $k$ ¿ 0

A similar approach is applied for modes $k > 0$, for which $\sigma_k = 0$. Eliminating the growth rate yields the following constraints on the mode amplitudes:

$$r_k=\frac{r_{\rm on}+\bar{r}+Dk^2}{\epsilon(1+\log\bar{r})/R_k-\bar{n}/n_k}, \tag{11a}$$

$$\bar{n}LR_k^2+\left[\left(1+\log\bar{r}+\zeta_1\frac{\bar{n}}{\bar{r}}\right)(r_{\rm on}+\bar{r}+Dk^2)-\epsilon(1+\log\bar{r})L-\zeta_1\bar{n}\log\bar{r}\right]n_kR_k+\epsilon\zeta_1\log\bar{r}(1+\log\bar{r})n_k^2=0, \tag{11b}$$

where $L = (h\bar{\sigma}_m/\bar{R}^2)(k^2-1)$. The corresponding growth rate satisfies the quadratic equation

$$\left(1+\log\bar{r}+\zeta_1\frac{\bar{n}}{\bar{r}}\right)\lambda_k^2+\left[\left(1+\log\bar{r}+\zeta_1\frac{\bar{n}}{\bar{r}}\right)(r_{\rm on}+\bar{r}+Dk^2/\bar{R}^2)+\frac{\epsilon h\bar{\sigma}_m}{\bar{R}^2}(1+\log\bar{r})(k^2-1)-\zeta_1\bar{n}\log\bar{r}\right]\lambda_k$$
$$+\frac{\epsilon h\bar{\sigma}_m}{\bar{R}^2}(1+\log\bar{r})(k^2-1)(r_{\rm on}+\bar{r}+Dk^2/\bar{R}^2)=0. \tag{12}$$

The properties of the solutions are different for $k = 1$ and $k > 1$, so we discuss these two cases separately. For the $k = 1$ mode, there is only one real eigenvalue

$$\lambda_1=\frac{\zeta_1\bar{n}\log\bar{r}}{1+\log\bar{r}+\zeta_1\bar{n}/\bar{r}}-(r_{\rm on}+\bar{r}+D/\bar{R}^2), \tag{13}$$

and this mode is unstable when $\lambda_1 < 0$, providing the following criterion:

$$\zeta_1 r_{\rm on}(\bar{r}\log\bar{r}-r_{\rm on}-\bar{r}-D/\bar{R}^2)<\bar{r}(r_{\rm on}+\bar{r})(r_{\rm on}+\bar{r}+D/\bar{R}^2)(1+\log\bar{r}). \tag{14}$$

For the $k > 1$ modes, the system is linearly stable when the real parts of the growth rates are both negative, namely,

$$\lambda_1\lambda_2=\frac{\epsilon h\bar{\sigma}_m(1+\log\bar{r})(k^2-1)(r_{\rm on}+\bar{r}+Dk^2)}{\bar{R}^2(1+\log\bar{r}+\zeta_1\frac{\bar{n}}{\bar{r}})}>0, \tag{15}$$

$$\lambda_1+\lambda_2<0. \tag{16}$$

The first condition always holds for $k > 1$, and the second condition gives the condition

$$\zeta_1 r_{\rm on}(\bar{r}\log\bar{r}-r_{\rm on}-\bar{r}-Dk^2)<\bar{r}(r_{\rm on}+\bar{r})(1+\log\bar{r})\left[r_{\rm on}+\bar{r}+Dk^2+\frac{\epsilon h\bar{\sigma}_m}{\bar{R}^2}(k^2-1)\right]. \tag{17}$$

#### 2 Geometry and related numerical methods: $\theta - L$ formation for an evolving planar curve

We parametrize the curve $\Gamma(t)$ by a Lagrangian parameter $\alpha \in [0, 2\pi]$. The geometry of our model can be categorized as a type of problem where the position vector $\boldsymbol{x}(\alpha, t)$ labeled by the Lagrangian tracer $\alpha$ evolves by a normal velocity $U$:

$$\frac{\partial \boldsymbol{x}}{\partial t}(\alpha, t) = U(\alpha, t)\boldsymbol{n}. \tag{18}$$

The arclength along the curve can be expressed as

$$s(\alpha, t) = \int_0^\alpha \left|\frac{\partial \boldsymbol{x}}{\partial \alpha'}(\alpha', t)\right| \mathrm{d}\alpha', \tag{19}$$

and the $s$ and $\alpha$ derivatives can be exchanged through

$$\partial_\alpha = s_\alpha \partial_s, \quad s_\alpha = |\boldsymbol{x}_\alpha| = \sqrt{x_\alpha^2 + y_\alpha^2}. \tag{20}$$

At each instant of time $t$, we can define the Frenet-Serret frame as the following:

$$\boldsymbol{t}(s) = \boldsymbol{x}_s, \quad \boldsymbol{n}(s) = -\frac{1}{\kappa}\boldsymbol{t}_s, \tag{21}$$

as sketched in Fig. 1($a$), where the curvature reads $\kappa = |\boldsymbol{x}_{ss}| = y_{ss}/x_s$ and

$$\boldsymbol{t}_s = -\kappa \boldsymbol{n}, \quad \boldsymbol{n}_s = \kappa \boldsymbol{t}. \tag{22}$$

Denoting the angle between the unit tangent vector and the $x$-axis as $\theta(\alpha, t)$, the unit tangent and normal vectors can also be expressed in terms of $\theta$ as

$$\boldsymbol{t} = \begin{pmatrix} \cos\theta \\ \sin\theta \end{pmatrix}, \quad \boldsymbol{n} = \begin{pmatrix} \sin\theta \\ -\cos\theta \end{pmatrix}, \tag{23}$$

and the curvature is $\kappa = \theta_s = \theta_\alpha / s_\alpha$. Given the tangent angle $\theta(\alpha, t)$ and the local arclength derivative $s_\alpha$, the shape of the curve can be reconstructed by

$$\boldsymbol{x}(s, t) = \boldsymbol{x}(0, t) + \int_0^s \boldsymbol{t}(s', t)\mathrm{d}s' = \boldsymbol{x}(0, t) + \int_0^\alpha \boldsymbol{t}(\alpha', t) s_{\alpha'} \mathrm{d}\alpha', \tag{24}$$

and $\theta$ and $s_\alpha$ can be chosen as independent dynamical variables instead of $\boldsymbol{x}$.

Since the shape of the curve evolves only by the given normal velocity $U$, an arbitrary tangential velocity $T$ can be introduced without changing the shape and only provides a change in frame for the parametrization [1]. After introducing the tangential velocity, the shape equation now becomes

$$\frac{\partial \tilde{\boldsymbol{x}}}{\partial t}(\alpha, t) = U\boldsymbol{n} + T\boldsymbol{t}. \tag{25}$$

Note that curves represented by $\tilde{\boldsymbol{x}}$ and $\boldsymbol{x}$ have the same shape, but with different positions of material points, see Fig. 1($b$). In the case of interfacial flows, we are free to apply this coordinate transform since only the interfacial geometry is the matter of interest. However, for an evolving material line, a tangential velocity will transport material points along the curve. As a consequence, the kinematic equation for physical quantities along the curve should be modified as

$$\frac{\partial \tilde{n}}{\partial t} = r_{\text{on}}(1-\tilde{n}) - \tilde{r}\tilde{n} + D\frac{\partial^2 \tilde{n}}{\partial \tilde{s}^2} + \frac{T}{\tilde{s}_\alpha}\frac{\partial \tilde{n}}{\partial \alpha}, \tag{26}$$

where $\tilde{n}$ denotes the value of $n$ at the new grid points. The shape evolution of the curve is determined by the $\theta$–$L$ formulation,

$$\frac{\mathrm{d}L(t)}{\mathrm{d}t} = \int_0^{2\pi} U\tilde{\theta}_\alpha \mathrm{d}\alpha, \tag{27}$$

$$\frac{\partial \tilde{\theta}}{\partial t}(\alpha, t) = -\frac{U_\alpha}{\tilde{s}_\alpha} + \frac{T\tilde{\theta}_\alpha}{\tilde{s}_\alpha}, \tag{28}$$

where $L(t)$ is the total length of the curve and the tangential velocity $T$ satisfies

$$T(\alpha, t) = -\int_0^\alpha \tilde{\theta}_\alpha(\alpha', t)U\mathrm{d}\alpha' + \frac{\alpha}{2\pi}L_t \tag{29}$$

to keep the mesh points equally spaced at each instant of time.

### 3 Numerical validation

We validate our numerical method by comparing the short time evolution of an isolated Fourier mode to predictions of the linear stability analysis. Results for $r$, $n$ and $R$ for more $k = 15$ are plotted in Fig. 2, where excellent agreement is observed in the linear regime.

### 4 Parameter estimation

Following [2], we list the estimated values of parameters in Table 1, where the various references used are provided.

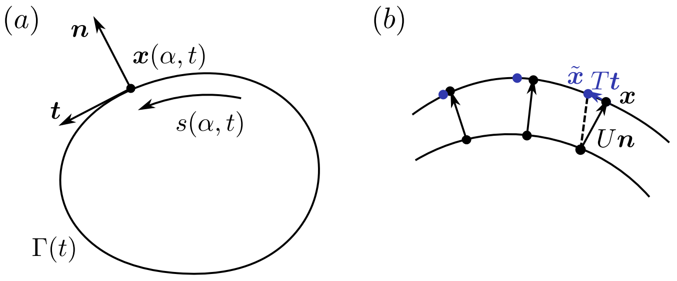


Figure 1: Geometry and parametrization: (a) Frenet-Serret frame. (b) Schematic description of the material points evolving without the normal velocity (black, $\boldsymbol{x}$) and with the tangential velocity (blue, $\tilde{\boldsymbol{x}}$)

.

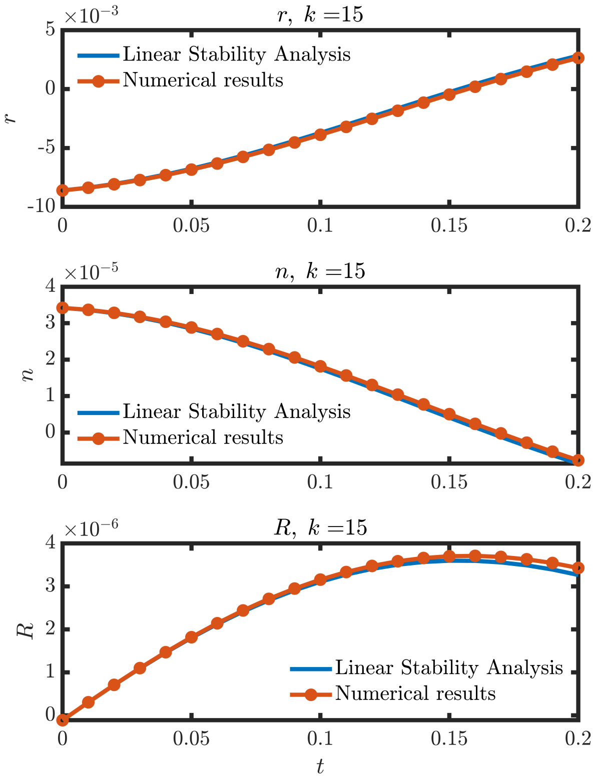


Figure 2: Temporal evolution of the off rate $r$, fraction of bound linkers $n$, and radius $R$ for an initial perturbation given by an isolated Fourier mode of wavenumber $k = 15$. The figure compares results from numerical simulations with predictions from the linear stability analysis.

Table 1: Parameters used in the model with sources.

| Parameter | Symbol | Estimation | Dimension |
|---|---|---|---|
| On-rate | $k_{\text{on}}$ | 1 [3] | $\text{s}^{-1}$ |
| Off-rate under zero force | $k^0_{\text{off}}$ | 0.1 [3] [4] | $\text{s}^{-1}$ |
| Typical rupture force | $f_0$ | 5 [5] [3] | pN |
| Spring stiffness | $k_b$ | 0.5 [2] [3] | pN/nm |
| Bond density | $\rho$ | $10^{-4} - 10^{-5}$ [2] [6] | $\text{nm}^{-2}$ |
| Diffusion coefficient of adhesions | $D$ | 0.2 [7] [8] | $\mu\text{m}^2/\text{s}$ |
| Viscous drag friction coefficient | $\zeta_0$ | 100 [2] | Pa·s |
| Stochastic friction coefficient | $\zeta_1$ | $10^5$ | Pa·s |
| Polymerization velocity | $v_p$ | 100-200 [9] [10] | nm/s |
| Cell radius | $R_0$ | 10 [11] | $\mu$m |
| Lamellipodium height | $2h$ | 140-200 [12] [13] | nm |
| Lamellipodium width | $l_1$ | 2-4 [11] [14] | $\mu$m |
| actomyosin contractile tension | $\sigma_c$ | 2 | pN/nm |
| Membrane area stretch modulus (fish keratocyte) | $k$ | $10^5$ [7] | pN/$\mu$m |
| Effective stiffness (2D) | $k_\sigma = k/R_0^2$ | $10^3$ | pN/$\mu\text{m}^3$ |


[7] Erin Lynn Barnhart. *Oscillations, waves, and symmetry breaking in cell motility.* PhD thesis, Stanford University, 2010.
